# Evolution of Learning in Technology Adoption: The case of the U.S. Soybean Seed Industry

**DOI:** 10.1101/2021.07.22.453433

**Authors:** Cornelia Ilin, Guanming Shi

## Abstract

This paper examines how the evolution of learning affects technology adoption. We use a sequential adoption model that accounts for differences between forward-looking adopters, who consider future impacts of their learning, and myopic adopters, who only consider past learning. We apply the analysis to three panels of U.S. soybean farmers representing different stages of the genetically modified (GM) seed technology diffusion path. We show that uncertainty is considerably reduced over time due to increased learning efficiency. Our results indicate that a forward-looking model fits the early adopters and early majority stages better, while both models perform equally well in the laggard stage.

**JEL classification:** D83, Q31, Q33, Q16

## Introduction

New technology diffusion and adoption are two sides of the same coin—with “diffusion” referring to the rate at which a new technology spreads among all potential users and “adoption” describing an individual user’s decision to try or not to experiment with new technology. Researchers in the fields of sociology, anthropology, education, social networks, and communications have extensively studied the technology diffusion path [1]. The S-shaped path – broken down into “early adopters”, “early majority”, “late majority”, and “laggard” stages – is commonly observed in many cases and is a well-known result of these studies. Following [1-2], “early adopters” and “early majority” are defined as the stages where the number of new adopters increases but with different trends: faster and slowly, respectively; and “late majority” and “laggards” are the stages where the number of new adopter decreases, also with the different trend: slowly and faster, respectively [1-5]. Economists, on the other hand, have studied adoption decisions—developing models that identify, examine, and characterize the various factors that may influence a user’s technology adoption decisions [6]. In this literature, binary choice models, wherein users choose to adopt a new technology or choose not to adopt [7-10], are distinguished from sequential adoption models, wherein users initially and partially adopt a technology, and then adjust adoption practices in later years [11,12]. Some researchers model myopic adoption behavior with immediate utility maximization only [13-15], while others model forward-looking behavior where adopters maximize utility over a planning horizon [16-18].

While most papers on adoption allow for time -varying covariates that may change over time, there is limited study documenting explicitly whether and how adoption decisions change in different stages of the technology diffusion path. Changes in adoption decisions may be particularly relevant when new technology adoption is associated with changes in user knowledge and learning after initial uncertainty about the new technology’s functionality. This uncertainty is often seen when potential users are heterogeneous and/or the technology will be used in heterogeneous situations. For example, in the case of agricultural technology such as seeds or pesticides, farmer knowledge and resources can vary widely while their crop plots may differ in soil quality and vulnerability to potential infestations. Farmers may adjust adoption decisions based on learning about profitability by observing neighbors’ experiences or from their own past experiences, which could differ by diffusion stage.

This study aims to understand how learning from adoption decisions evolves in different stages of the diffusion path. It may provide important guidance to researchers on how and when to choose the appropriate static or dynamic model in analyzing adoption decisions. For example, if there are not many learning dynamics at a certain stage of the adoption path, then the simple static model would be appropriate without loss of insight into potential adopters’ learning. Moreover, the information on how farmers’ learning evolves is also helpful for policymakers and marketing companies in designing effective strategies promoting new technology adoption.

In particular, we examine U.S. farmers’ adoption of new genetically modified (GM) soybean seed technology from its introduction in 1996 to 2009. As Fig 1 suggests, GM soybean seed technology exhibits a completed diffusion path: from 2 percent adoption in 1996 to 75 percent in 2001, then 95 percent adoption in 2009 where it currently remains.

**Fig 1.**
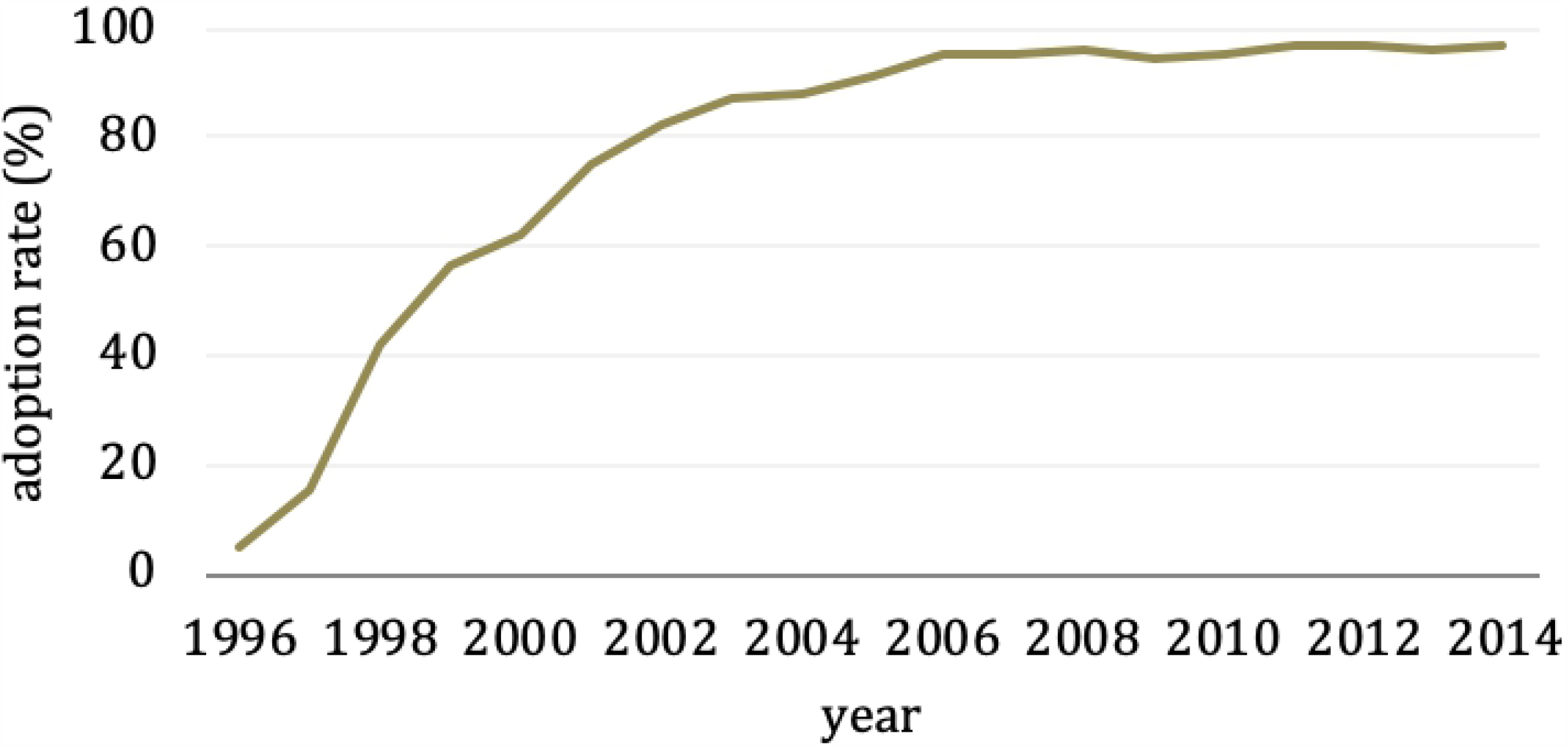
Adoption rate of GM soybean seeds in the U.S.: Year-level comparison of adoption rates of GM soybean seeds between 1996 to 2014. The data used is farm-level soybean seed purchase information collected by dmrkynetec, St. Louis, MO (dmrk).

To understand GM soybean seed adoption decisions, our work joins a growing literature that examines behavioral explanations for sequential adoption of new technologies under uncertainty and learning [11,19,20-22]. Following previous studies, we hypothesize two sequential models with Bayesian learning in which farmers initially update believes about GM seed profitability after observing noisy signals from their own and neighbors’ experience. We follow [22] to specify and estimate (i) a “myopic” model in which farmers adopt technology at a rate that provides the highest current expected utility, and (ii) a “forward-looking” model in which farmers recognize that current technology choices will affect their future information sets.

However, our work extends the modeling and analysis presented in [22]. By looking at farmers’ adoption decisions in different stages of the diffusion path, we obtain insights about the evolution of learning and uncertainty. We apply our analysis to three distinct panels of soybean farmers in the U.S. for three time periods: 1996 – 2001, 2001 – 2006, and 2006 – 2009. We did not include the 2010 – 2014 time period, because the market reached saturation starting in 2010. Following [3,4]’s definition of different diffusion stages, we label the 1996 – 2001 period the “early majority” stage, the 2001 – 2006 period the “late majority” stage, and the 2006 – 2009 period the “laggard” stage. Concretely, we fit a polynomial specification to the adoption path and determine the stage categories by examining the sign of the first and second-order of the rate of changes in adoption increments. These periods correspond to adoption rates changing from zero adoption to 75%, then from 75% to 95%, and finally stable at around 95%. Our results suggest that uncertainty about profit reduces significantly after farmers experiment with the new seed technology for a number of years. This decrease is attributed to less noisy signals from their own and neighbors’ experiences, which, in turn, is associated with an increase in learning efficiency. We also find that the “forward-looking” sequential model outperforms the “myopic” model in generating the observed adoption path for the 1996-2001 and 2001-2006 farmer panels, while both models perform equally well for the 2006-2009 farmer panel. This may be because diffusion is close to saturation beginning in 2006, thus little uncertainty remains about the new technology’s performance or attributes.

## Methods

### Learning models

In this section, we describe the “myopic” and “forward-looking” sequential learning models following [22] in the context of GM soybean seed technology adoption. Detailed derivations can be found in [22].

Consider farmer *i*, who decides in each time period *t* = 1,…,*T* which seed technologies to plant on her farmland of size *A*_*i*_ . She has two choices: an existing conventional technology (old) and/or a newly developed GM technology (new). Let *Git*_*it*_ denote the amount of farmland planted with the new technology by farmer *i* at time *t*. Farmer *i* chooses the optimal adoption rate of the new technology 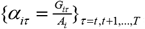, to maximize the expected present utility value over the planning horizon. The value function of farmer *i* at time *t, V_it_* (·), is defined as:

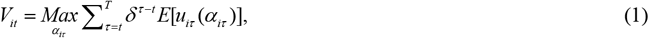

where *δ* > 0 is the discount factor and *u*_*iτ*_ is the utility of farmer *i* at time *τ*.

The Bellman equation is:

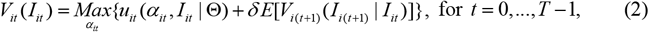

where *I*_*it*_ is the information set of farmer *i* at time *t*, and Θ is a set representing the parameter space to be defined in the model. Equation (2) indicates that the value of choosing a specific adoption rate at time t equals the sum between the immediate utility of choosing *αit* (i.e. *u*_*it*_ (*α*_*it*_, *I*_*it*_ | Θ)) and the additional expected value of choosing *α*_*it*_ of having the augmented information set *I*_*i(t +* 1)_ (i.e. *δ*E [*V*_*i(t +* 1)_ (*I*_*i(t +* 1)_ | *I*_*it*_)]). This specification provides an information gathering incentive. For example, a farmer may choose to partially adopt the new technology, even if doing so provides a lower expected utility than with zero adoption, as long as this decision leads to a sufficient increase in the value of information she will have in the following time period. Therefore, we define farmers with immediate expected utility maximizers (where *δ* = 0) the “myopic” ones, and those with expected utility maximizers over a planning horizon (where *δ* > 0) the “forward-looking” ones.

We assume that the distribution functions of profits for both seed technologies are unknown to farmers. Following [22], we construct farmer *i*’s expected utility at time *t* as a function of the mean and variance of the total profits from planting the two types of seeds: *πit* = *π* _*igt*_ + *π* _*ict*_, where the subscript “*g*” denotes the GM seed type, and the subscript “*c*” denotes the conventional seed type. Farmer learning about GM seed profitability, and hence the adoption process for both myopic and forward-looking farmers, can be developed accordingly (refer to the appendix for details).

The current payoff at time t for farmer i is:

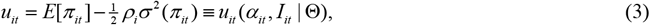

where

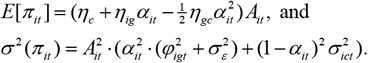

The derivation of the mean (*E*[*π*_*it*_]) and variance of the total profits (*σ*^2^ (*π*_*it*_) can be found in the appendix, or in [22]. As defined earlier, *α*_*it*_ is farmer *i*’s adoption rate of the GM seed technology in her farmland with total acreage *A*_*it*_ at time *t*. We use *ρi* to measure farmer i’s degree of risk aversion, which is assumed to be inversely related to the acreage *A*_*it*_ . The parameter *η*_*c*_ represents the mean profit of planting conventional seed per plot, which is common to all farmers, while *η*_*ig*_ is the upper bound of the plot level profit difference between planting GM seed and conventional seed. Additionally, *η*_*gc*_ measures the rate of dissipation of such profit difference along with adoption. Factors affecting the variance of the total profits include adoption rate *α*_*it*_, total acreage *A*_*it*_, the current belief of GM profit variance 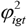, the profit variance of conventional seeds 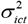 and the time invariant GM profit variance 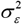 .

Equation (3) suggests that farmer’s utility function is determined by the choice variable (adoption rate *α* _*it*_) and the information set *I*_*it*_, given the remaining parameter space Θ . In our model, the information set *I*_*it*_ consists of factors that affect current expected utilities and/or the probability distribution of the future expected utilities (i.e. 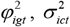 and *A*_*it*_). The parameter space set Θ include parameters used to define the variance of conventional seed profitability, the risk aversion measurement, the mean profits of planting conventional seeds and the mean profit differences between conventional seeds and GM seeds, the time invariant GM profit variance 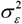 (which can be obtained by learning from own experience), a parameter measuring the variance in learning of GM profitability from neighbors 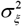, and the white noise added to the variance of the conventional seed profitability 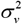.

### Data

For our empirical analysis, we use U.S. farm-level soybean seed purchase data collected by dmrkynetec, St. Louis, MO (dmrk). Data are obtained from annual surveys on a stratified sample of U.S. soybean growers since the introduction of GM soybean seed in the U.S. in 1996. It includes information on seed type, quantity, and prices, and on farm size for 26,314 farmers between 1996 and 2014. To capture the evolution of uncertainty and learning, we examine farmer’s adoption decisions in three distinct stages along the diffusion path: early majority (1996 – 2001), late majority (2001 – 2006), and laggard (2006 – 2009). Note that we do not examine 2010 – 2014 as the adoption rate seems to have reached saturation limit by 2010 (Fig 1). We chose farmers who farmed soybeans in each year of the time periods of each of our three study panels. We further eliminate farmers with total soybean acreage changing dramatically over the sample period to remove those with frequent switching behavior between soybean and other crops. We do this by constructing a farm acreage variation measure by dividing the standard deviation of the farm acreage by its mean. We dropped those farmers with greater than 40 percent variation. We also dropped Crop Reporting Districts (CRD) (as defined by the United States Department of Agriculture (USDA)) in which the number of surveyed farmers fell below the 10 percent quantile (the threshold number of farmers is 8 in 1996 – 2001, 11 in 2001 – 2006, and 8 in 2006 – 2009) to ensure statistical significance in calculating CRD-level market characteristics. Our panel data contain 377 early majority adopters, 190 late majority adopters, and 473 laggard adopters. Fig 2 shows the average farmer adoption rate of GM soybean seeds from our three study panels (1996-2009), from the whole dmrk data (1996-2009), and USDA data (2000 to 2009, based on data availability) [26]. All adoption data follow a similar pattern, suggesting that farmers in our samples do not differ from those in the population at large in terms of adoption behavior over time.

**Fig 2:**
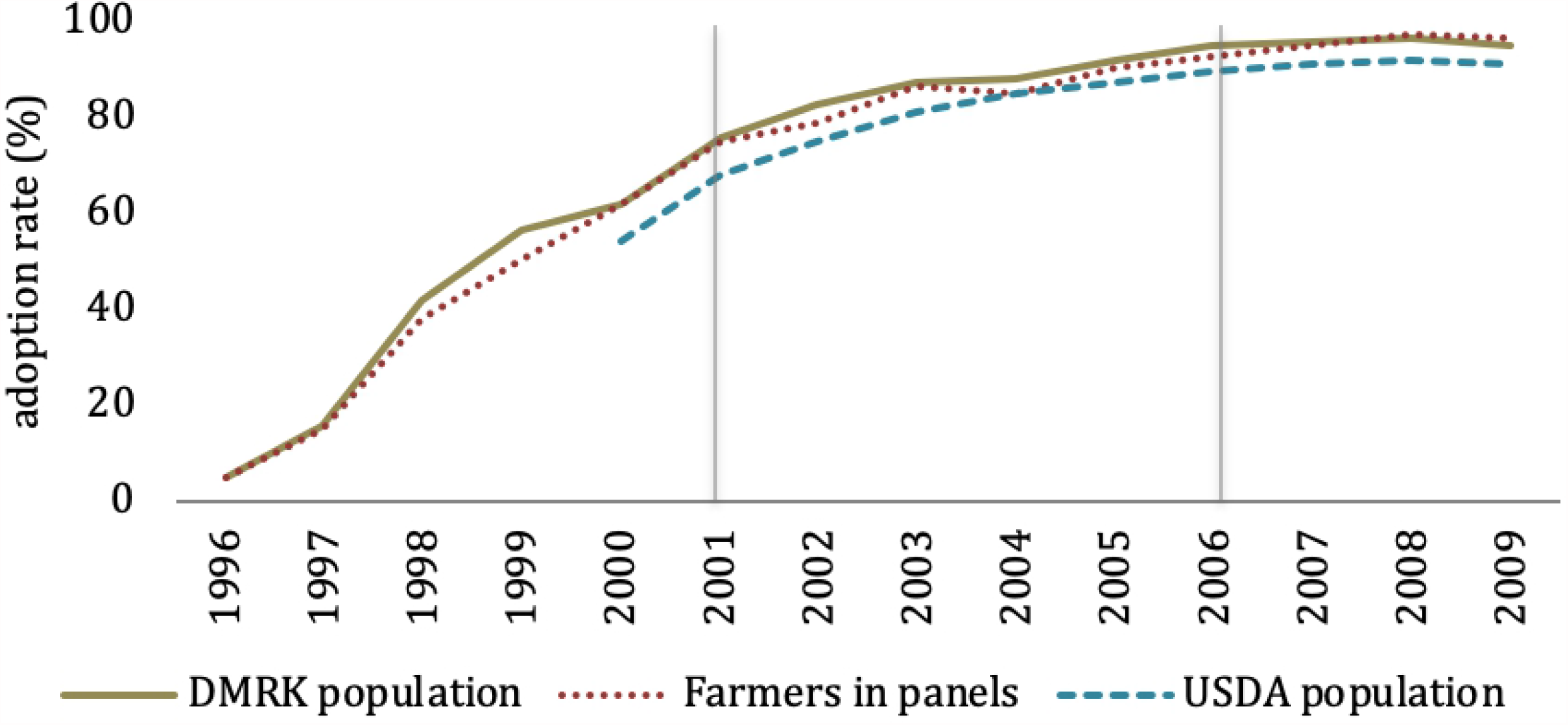
Average adoption rate of GM soybean seeds in the U.S.: sample vs. population. Comparison of DMRK population (khaki long line) vs. farmers in panels (red dotted line) vs. USDA population (blue dashed line) adoption rate of genetically modified soybean seeds in the U.S. From left to right, the first vertical solid grey line indicates the end of the “early majority” stage (1996 – 2001) and the second line indicates the end of the “late majority” stage (2001 – 2006).

We believe learning from neighbors is valuable if neighbors face similar agro-climatic conditions. Therefore, we use CRD as a proxy for local market. This aligns with common practice in the empirical literature, in which the neighborhood effect is based on geographical proximity [11,23,24]. We construct the CRD adoption rate using the dmrk population data. Thus, while *A*_*it*_ is measured by individual farmer’s total soybean acreage in year *t, A*_–*it*_is calculated by the average soybean acreage for other farmers in the CRD in that year. We also include both linear and quadratic forms of latitude and longitude of the center of the county where each farmer is located. This captures spatial heterogeneity in farming systems and agro-climatic conditions (e.g. temperature, rainfall volume, and daylight length).

Table 1 shows summary statistics of the variables for each panel of farmers. The average soybean acreage for farmers in each panel is notably larger than that of their neighbors. It seems likely that DMRK oversamples big farms, and that big farms are more likely to stay in the panel than small ones. Yet Fig 2 shows that there may not be much attrition bias for our study purpose. Descriptive statistics also suggest that panel farmers and their neighbors pay similar prices for conventional and GM seeds. Also, seed prices increased considerably between 1996-2001 and 2006-2009. The average price premium for GM seeds relative to conventional also increased from around $8.5 in 1996-2001, to about $13 in 2006-2009; yet, price difference between conventional and GM stays at around 85 percent. Latitude and longitude variables show that farmers in the three panels are located in the U. S. Midwest.

**Table 1.** Variable description and summary statistics.

| Variable | Description | 1996-2001 |  | 2001-2006 |  | 2006-2009 |  |
| --- | --- | --- | --- | --- | --- | --- | --- |
|  |  | Mean | SD | Mean | SD | Mean | SD |
| $\alpha_{it}$ | Farmer $i$'s adoption rate at time $t$ | 0.41 | 0.45 | 0.84 | 0.34 | 0.96 | 0.19 |
| $\alpha_{it-1}$ | Farmer $i$'s adoption rate at time $t-1$ | 0.34 | 0.42 | 0.82 | 0.35 | 0.96 | 0.19 |
| $\alpha_{-it}$ | Farmer $i$'s neighbors' adoption rate at time $t$ | 0.43 | 0.29 | 0.86 | 0.11 | 0.95 | 0.1 |
| $\alpha_{-it-1}$ | Farmer $i$'s neighbors' adoption rate at time $t-1$ | 0.37 | 0.27 | 0.84 | 0.11 | 0.96 | 0.1 |
| $A_i$ | Farmer $i$'s total soybean acreages | 258 | 255 | 305 | 310 | 398 | 398 |
| $A_{-i}$ | Farmer $i$'s neighbors' total soybean acreage | 95 | 39 | 97 | 44 | 118 | 94 |
| $P_{it}^{GM}$ | GM seed price paid by farmer $i$ at time $t$ (\$/50 lb. bag) | 19.71 | 4.06 | 23.78 | 3.73 | 31.64 | 5.99 |
| $P_{-it}^{GM}$ | GM seed price paid by farmer $i$'s neighbors at time $t$ (\$/50 lb. bag) | 19.53 | 3.57 | 23.59 | 2.92 | 31.54 | 5.34 |
| $P_{it}^{Conv}$ | Conventional seed price paid by farmer $i$ at time $t$ (\$/50 lb. bag) | 11.37 | 4.03 | 12.69 | 4.35 | 18.09 | 4.68 |
| $P_{-it}^{Conv}$ | Conventional seed price paid by farmer $i$'s neighbors at time $t$ (\$/50 lb. bag) | 10.61 | 2.82 | 12.31 | 4.26 | 18.08 | 4.68 |
| Lat | Latitude of the farm | 41.05 | 2.33 | 41.33 | 2.35 | 41.15 | 2.8 |
| Lon | Longitude of the farm | 89.71 | 5.05 | 91.08 | 4.62 | 90.81 | 4.99 |

Description of variables and summary statistics (mean and standard deviation (SD)) of variables used in the analysis for “early majority” (1996-2001), “late majority” (2001-2006), and “laggard” (2006-2009) samples of farmers.

### Estimation

In the empirical application, we apply the structural models discussed in the Methods-Learning Models section to each panel of farmers described in Methods-Data section. For each model, we use the Nelder-Mead simplex method to minimize the simulated generalized method of moments (GMM) objective function. More precisely, we search for the set of parameters that minimize a weighted distance between the optimal (predicted) adoption path and the observed adoption path (see [22] for details). The instruments used to facilitate the estimation are presented in the next section.

### The “myopic” model

Myopic farmers choose the adoption rate that gives the highest current period expected utility. They maximize the value function defined in equation (1), with discount factor set to *δ* = 0 and utility function as specified in equation (3):

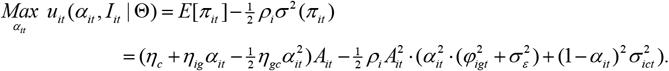

The first order condition gives:

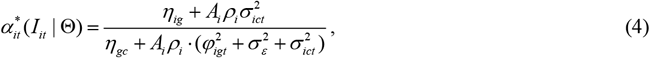

where Θ is the parameter space as defined before; and the second-order condition holds.

To compute the predicted adopted path for each farmer in each panel following equation (4) we need to update the Bayesian beliefs on the variance of GM seed mean profitability 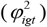 after each time period. Because the actual adoption rate in the first year of each panel (1996, 2001, 2006) is known (noting that that observations form the first year of each time period sample are used as a benchmark only), we can obtain the perceived GM variance for the first period 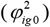 by solving the inverse function of 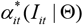 for *t* = 0 . We then update 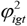 for all the following years according to the Bayesian rule in equation (A2) in the appendix.

### The “forward-looking” model

Forward-looking farmers choose the adoption rate that maximizes the expected present value of utility over a planning horizon. They perceive that their current choices affect their information sets, and this perception creates an incentive to experiment to learn about the technology. The optimization problem follows the value function defined in equation (1), with discount factor *δ* > 0 and utility function defined in equation (3):

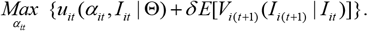

To compute the predicted adoption path, we make the following assumptions:

#### The transition probabilities

We rewrite *A*_*it*_ and *A*_*−it*_ as *A*_*i*_ and *A*_*−i*_ because we focus on farmers with relatively stable farm size over time. Thus, the information set can be reduced to 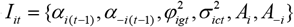 Following [11] we write the transition rules for all these variables in Markovian form (see [22], section 4.2.1 for details).

#### The value function of the last period

Data suggest that toward the end of 2001 the change in adoption rate decreased in slope and flattened after 2006 (see Fig1). As such, we assume that in the last period of each sample, sequential learning reaches steady state, *E (V*_*i*_*(I*_*t+1*_)) = *E(V*_*i*_*(I*_*t*_*)) t* ≥ 6 in the 1996-2001 and 2001-2006 panels, and for *t* ≥ 4 in the 2006-2009 panels.

#### The prior for the first period Bayesian beliefs

The relationship between Bayesian beliefs and farmers’ adoption rates is no longer a one-to-one correspondence as in the “myopic” model; therefore, we cannot obtain first period priors, 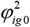, from the first year’s actual adoption rates. To overcome this problem we use beliefs of each farmer in year 1996, 2001, and 2006 in the myopic case as reference values in the forward-looking case. We add a parameter *b* to all these myopic beliefs to account for the potential bias introduced by the assumption. For example, the perceived variance of GM seed mean profitability in year 1997 for farmer i in panel 1996-2001 follows the Bayesian rule:

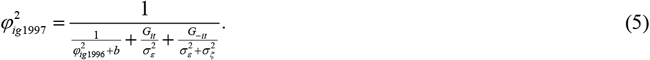

The same rule is applied for year 2001 in panel 2001-2006 and year 2006 in panel 2006-2009.

## Results

In estimation of the “forward-looking” model we chose the discount factor at *δ* = 0.96 . In addition to the discount factor *δ* and the additional bias parameter *b*, there are another 14 parameters to be estimated as listed in Table 2. These 14 parameters are shared by the “myopic” model. The initial values for the “myopic” model for each panel of farmers follow [22] (see Table 2). We use the estimated parameters in the “myopic” model as starting values for the “forward-looking” model.

**Table 2.** Initial values for parameters of the “myopic” model.

| Parameter | Definition | Initial value |
| --- | --- | --- |
| $\eta_{ig}$ | Constant term of GM mean profit | 1.00 |
| $\eta_{gc}$ | Decreasing rate of GM mean profit with adoption | 0.50 |
| $\gamma_0$ | Constant term for Conv variance | 5.00 |
| $\sigma_\epsilon^2$ | GM profit variance | 1.00 |
| $\sigma_\xi^2$ | Learning variance from neighbors | 10.00 |
| $\beta_0$ | Risk averse | 1.00 |
| $\beta_1$ | Farm size effect on risk averse | 0 |
| $\lambda$ | Parameter of AR1 process | 0.20 |
| $\sigma_v^2$ | Disturbance of AR1 process | 0.24 |
| $\gamma_1$ | Linear term for Conventional variance | 1.00 |
| $c1_{lat}$ | Latitude effect on mean profit | -0.19 |
| $c2_{lat2}$ | Second-order effect of latitude | 0.08E-02 |
| $c3_{lon}$ | Longitude effect on mean profit | 0.25 |
| $c4_{lon2}$ | Second-order effect of longitude | -0.03E-02 |
| $b$ | Adjustment on Bayesian beliefs on GM variance | 0 |

The initial parameter values for the “myopic” model for each panel of farmers: “early majority” (1996-2001), “late majority” (2001-2006), and “laggard” stage (2006-2009).

We choose 17 instruments to estimate GMM: a constant vector 1, total soybean acreage of farmer *i* and his neighbors (*A*_*i*_, *A*_*−1*_) plus the square terms, previous year adoption rate (*α* _*i(t* − 1)_, *α* _−(*t* −1)_) and the square terms, farm characteristics **X**_*i*_ (latitude and longitude of each county center where farms are located and the square terms). Although not explicitly included in our model, pricing also affects adoption decisions. As such, we add conventional and GM seed prices paid by farmer i and his neighbors (computed as the average price in a given CRD).

Parameter estimates and standard errors for both “myopic” and “forward-looking” models are presented in Table 3. Our results suggest that the “forward-looking” model predicts a different adoption scenario than the “myopic” model for the 1996-2001 panel of farmers. Differences in predictions between the two models significantly decrease for the 2001-2006 panel. For the 2006-2009 panel the predictions are very similar. One possible explanation is that, even if farmers are “forward-looking,” the value of the information gathering process declines. As farmers experiment with the technology their expectations for new leaning may also decrease.

**Table 3.** Estimation results for the “myopic” and “forward-looking” models.

| Parameter | Early Majority<br>(1996-2001) |  | Late Majority<br>(2001-2006) |  | Laggard<br>(2006-2009) |  |
| --- | --- | --- | --- | --- | --- | --- |
|  | “Myopic” | “Forward-looking” | “Myopic” | “Forward-looking” | “Myopic” | “Forward-looking” |
| $\eta_{ig}$ | 1.82***<br>(0.01) | 2.04***<br>(0.22) | 1.51***<br>(0.40) | 1.73***<br>(0.19) | 0.56***<br>(0.02) | 0.71<br>(1.07) |
| $\eta_{gc}$ | 0.42***<br>(0.01) | 0.42<br>(0.50) | 0.57***<br>(0.15) | 0.61<br>(0.46) | 0.57***<br>(0.06) | 0.24<br>(0.24) |
| $\gamma_0$ | 3.15***<br>(0.01e-2) | 1.42***<br>(0.26) | 6.09<br>(11.42) | 3.60***<br>(0.54) | 5.02<br>(74.71) | 4.31***<br>(0.98) |
| $\sigma_g^2$ | 0.83***<br>(0.03) | 0.82***<br>(0.04) | 0.34<br>(1.23) | 0.25**<br>(0.10) | 0.03<br>(0.43) | 0.03<br>(0.02) |
| $\sigma_\xi^2$ | 31.74*<br>(15.07) | 59.57<br>(39.59) | 18.41<br>(49.52) | 41.61***<br>(5.63) | 35.79<br>(515.75) | 96.04<br>(78.17) |
| $\beta_0$ | 1.19***<br>(0.06e-1) | 1.08***<br>(0.25) | 1.40<br>(4.39) | 1.30***<br>(0.12) | 1.09<br>(14.14) | 2.57<br>(4.05) |
| $\beta_1$ | -0.84***<br>(0.00) | -0.90***<br>(0.04) | -0.13<br>(0.93) | -0.14***<br>(0.03) | 0.05<br>(1.19) | -0.16<br>(0.14) |
| $\lambda$ | 0.21<br>(0.70) | 0.23***<br>(0.01) | 0.20<br>(13.49) | 0.25***<br>(0.05) | 0.23<br>(34.99) | 0.29<br>(0.71) |
| $\sigma_v^2$ | 0.46<br>(0.52) | 0.52<br>(0.32) | 0.32<br>(27.77) | 0.30*<br>(0.17) | 0.40<br>(46.51) | 0.37<br>(3.36) |
| $\gamma_1$ | 1.95<br>(2.26) | 1.81***<br>(0.57) | 0.87<br>(143.80) | 1.01<br>(3.20) | 0.53<br>(108.97) | 1.55<br>(8.03) |
| $c1_{lat}$ | 0.05***<br>(0.01e-1) | 0.05*<br>(0.02) | -0.18<br>(1.04) | -0.13<br>(0.33) | -0.11<br>(0.18) | -0.33<br>(0.40) |
| $c2_{lat2}$ | 0.05e-2***<br>(0.01e-2) | 0.06e-2<br>(0.05e-1) | 0.01e-1<br>(0.10) | 0.01e-1<br>(0.02e-1) | 0.09e-2<br>(0.07e-1) | -0.01e-1<br>(0.01e-1) |
| $c3_{lon}$ | 0.53***<br>(0.02e-1) | 0.53<br>(1.63) | 0.28<br>(2.10) | 0.32<br>(0.89) | 0.48***<br>(0.09e-1) | 0.17<br>(0.13) |
| $c4_{lon2}$ | -0.01e-2<br>(0.00) | -0.01e-2<br>(0.03) | -0.04e-2<br>(0.71) | -0.04e-2<br>(0.04e-2) | -0.05e-2<br>(0.13) | 0.22<br>(0.28) |
| $b$ | | 0.55***<br>(0.17) | | 0.18***<br>(0.01) | | 0.10<br>(0.13) |
| **Mean Square Error** | **2.485** | **0.177** | **0.185** | **0.049** | **0.012** | **0.011** |

Parameter estimates and standard errors (numbers in parenthesis) for the “myopic” and “forward-looking” models for each panel of farmers: “early majority” (1996-2001), “late majority” (2001-2006), and “laggard” (2006-2009).

Average squared root prediction errors for each particular year and each model (myopic. vs. forward-looking) are presented in Fig 3. In the following we present in more detail our findings for each panel of farmers.

**Fig 3.**
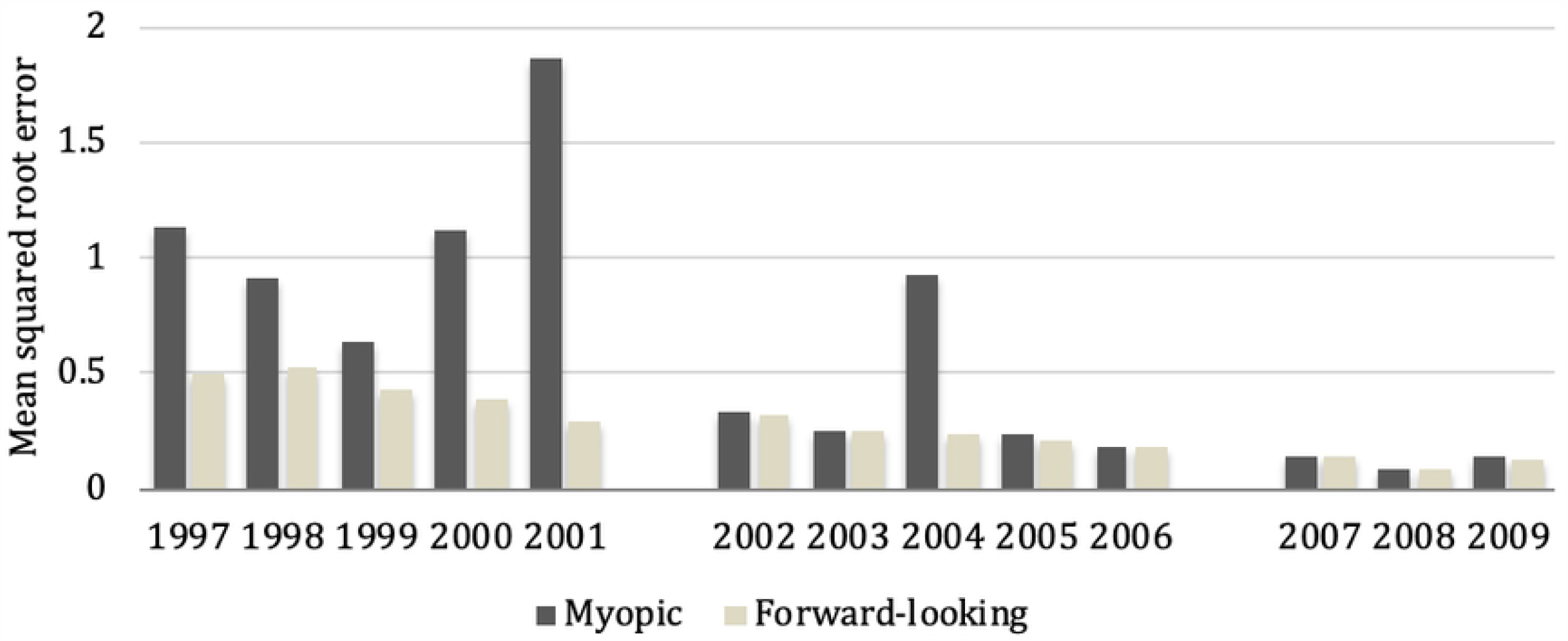
Mean squared root prediction errors for the “myopic” and “forward-looking” models. Year-by-year comparison of predictions in the “myopic” vs. “forward-looking” models across the three adoption stages: “early majority” (1996-2001, 2001-2006, 2006-2009).

### Model fit

For the “early majority” (1996-2001) and the “late majority” (2001-2006), results in Table 3 indicate that the mean square errors from the “myopic” model are larger (and more so for the early majority) than from the “forward-looking” model, suggesting that both early majority and late majority behave in the forward-looking way. Using the average root squared prediction error for each particular year, Fig 3 indicates that the “forward-looking” model predicts more accurately in each year from 1996 to 2001. On average, the actual adoption rate is 40.15% and the predicted myopic and forward-looking adoption rates are 48.84% and 41.81%, respectively. The “myopic” model predicts higher adoption rates in the first 6 years of experimenting with the new technology, suggesting that the myopic model may ignore the potential cost associated with adoption, thus lead to over-adoption.

However, along with learning in the very early stage of technology diffusion, farmers tend to shift from considering the future cost of adoption to incorporating future value of adoption in the second stage of diffusion. Thus, for the “late majority”, the myopic model and forward-looking adoption-rate projections move closer. The actual mean adoption rate is 83.63% and the predicted myopic and forward-looking adoption rates are 87.12% and 85.78%, respectively.

The difference disappeared completely in the laggard stage (2006 – 2009). Table 3 shows that both the “myopic” and “forward-looking” models fit the data equally well. The mean square errors of the two models are close to each other. This convergence may result from a completed learning process (farmers fully understand the quality or attributes of the new GM seed) as the new technology diffusion has reached saturation. The actual mean adoption rate is 96.90%, and this compares with the rate predicted by the “myopic” model of 96.5%, and by the “forward-looking” model of 95.75%.

The parameter *b* (Table 3) accounts for the difference in the Bayesian belief towards profit variance of GM seeds between myopic and forward-looking farmers. The parameter’s estimated value is positive and statistically significant for the “early majority” and “late majority” but not statistically significant for the “laggard” stage. This result suggests that the uncertainty about profitability of the new technology plays an important role in the first two stages along the diffusion path, yet with a declining magnitude.

### Self-learning versus learning from neighbors

Learning efficiencies, from own and neighbors’ experience, demonstrate different patterns along the diffusion path. We follow [11,22] to define a farmer’s own-learning efficiency from planting GM seeds on plot *K*_*i*_ as 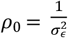, and the learning efficiency from his neighbor experience as 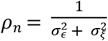

For the “early majority” stage (1996-2001), Table 3 shows that the parameter 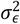 is positive and statistically significant, while the parameter 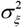 is not significant. These results indicate noise in learning from own experience but no additional learning noise from neighbor experience. Learning efficiencies from a farmer’s own experience and from his neighbors are about the same for the early majority (*ρ* _0_ = *ρ* _*n*_ 1.21).

For the “late majority” stage (2001-2006), however, the estimated parameters 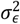 and 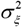 are both positive and statistically significant. Noise occurring in learning from neighbors is much louder than noise in self-learning. Consequently, the learning efficiency from a farmer’s own experience is much greater than that from neighbors’ experiences (*ρ* _0_ = 4 vs. *ρ* _*n*_ = 0.02)

For the “laggard” stage (2006-2009), the estimated parameters for 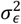 and 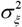 are not statistically significant. Consequently, there is perfect learning efficiency (*ρ* _0_ → ∞ and *ρ* _*n*_ → ∞). As we argued earlier, there might be nothing new to learn about the technology when diffusion is saturated.

Comparing learning efficiency patterns along the three stages on the diffusion path, it seems that farmers are cautious in the very early stage of the technology diffusion path and try to learn as much as possible from all possible sources, either own experiences or neighbors’ experiences. When knowledge is accumulated, farmers tend to count more on own experiences than on neighbors’ experiences. And when adoption reaches its saturation, learning seems to be complete from both sources. Note that [22] estimated soybean farmers’ adoption from 2001 to 2004, which is part of our so-defined “late majority” stage. They also found that farmers have higher learning efficiency from own experience than from neighbors’ experiences. Our analysis provides a more complete picture of the evolution of learning along the diffusion path.

### Mean profit

The estimated parameter *η*_*ig*_ represents the upper bound of the profit difference between conventional and GM technologies. It is positive, suggesting potential benefits of GM technology, yet with a decreasing magnitude moving along the diffusion path. And it is not statistically significant for the laggard stage. The *η*_*gc*_ parameter is not statistically significant for all three stages on the diffusion path, implying that the marginal profit from adopting GM seeds does not change when farmers allocate more acreage to GM seeds. The net benefits from adopting GM seeds, 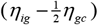 decrease along the diffusion path and approach zero at the end stage. It seems that when adoption increases, farmers successfully allocate lands so that those suitable for GM seeds and those suitable for the conventional seeds are planted with the corresponding appropriate technologies. The estimated parameters related to farm location (*c*1_*lat*_, *c*2_*lat*2_, *c*3_*lon*_, *c*4_*lon2*_) suggest that GM technology may have advantage over conventional seeds in the southern area for early majority adopters, but the advantage seems to disappear in the later adoption stages. Warm weather, tends to cause heavier weed infestation in southern areas, suggesting that early majority adopters are responding to severe weed infestation problems.

### Other results

For early to mid-stage adopters (1996 – 2001 and 2001 – 2006), the estimated risk averse parameters *β*_0_ are positive and significant, suggesting that early majority and late majority farmers in our samples are risk averse. They are also more risk averse if their farm size is larger, as the estimated parameter *β*_1_ is negative and significant. The estimated parameters *β*_0_, *β*_1_ are not statistically significant for model fitted for the “laggard” stage, indicating that farmers in our sample are not risk averse after 12 years of experimenting with GM soybean seed technology.

The effects of the random state variable on the variance of conventional seed profits (*γ*_1_) is positive and significant for the early majority (1996-2001), indicating that these farmers perceive a higher variance about profitability of conventional seeds. However, such patterns disappear towards the mid-and end-stage of the diffusion path, probably because farmers plant GM seeds on most of their land at this stage.

## Discussion

In this paper, we document the evolution of learning and uncertainty in the adoption of GM soybean seeds by farmers in the U.S. by examining farmers’ adoption decisions in three stages along the diffusion path: the “early majority” (1996 – 2001), the “late majority” (2001 – 2006), and the “laggard” stages (2006 – 2009). Unique data divided into three distinct panels of farmers surveyed annually between 1996-2001, 2001-2006, and 2006-2009, allows us to compare the results of a “forward-looking” model with a “myopic” model by modeling farmers’ adoption decisions in early-, midterm-, and last-stage of the diffusion path.

We find that the “forward-looking” model fits our data better than the myopic model in the “early majority” and “late majority” stages of the adoption process, suggesting that farmers in our samples are more likely to be forward-looking in the first 12 years of experimentation with the technology. Both models perform equally well in the last adoption stage, likely due to no differences in learning between myopic and forward-looking farmers once technology adoption has reached steady state.

We also find that farmers learn both from their own and neighbors’ experiences in the “early majority” and “late majority” stages, although neighborhood effect is considerably smaller in the late majority stage. In the “laggard” stage, learning is complete, resulting in minimal uncertainty from both sources. As a result, learning efficiency from own-experience and neighbor’s experience improves each year, resulting in a decrease in uncertainty about profitability of GM soybean seeds over time.

It is important to recognize farmers’ forward-looking behavior in “early majority” and “late majority” stages of the adoption process, and the similarity between myopic and forward-looking behavior in the “laggard” stage of the adoption process. For researchers studying technology adoption (or demand analysis of new products), our study provides guidance on how and when to choose the appropriate static or dynamic model. For early stages along the diffusion path, the dynamic model with future learning incorporated may provide a more accurate illustration of the market and demand. On the other hand, for a product in its later stages of the diffusion path, the simple static model would be appealing without loss of much insight in potential adopters’ learning. Another important finding is how farmers’ learning evolves over time. The fact that they switch from counting on neighbor learning to self-learning when moving along the diffusion path can serve as a source of information for policy makers or marketing firms interested in promoting the adoption of new agricultural technologies and could be extended to other market analyses. For example, it may be more effective to focus on providing training and extension support when introducing a new technology to farmers in the very early stage of the technology, and then focus more on subsidizing farmer adoption when the technology has already been in the market and adopted by certain percentage of potential users.

## Appendix

### Farmer expected utility

We assume farmers are uncertain about seed profitability for both technologies, i.e. the distribution function of profits is unknown. Let the total profit of farmer *i* from planting the two types of seeds in time *t* defined as: *π*_*it*_ = *π*_*ict*_ + *π*_*igt*_, where *π*_*ict*_ and *π*_*igt*_ are the profits from planting respectively conventional and GM seeds. Further, let the farmer’s expected utility be a function of the mean and variance of the total profit [25]:

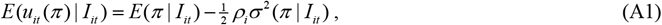

Where *E (π)* and *σ* ^2^ (*π*) are the mean and variance of the total profit, and *ρ* is a measure of farmer i’s degree of risk aversion, such that, following [22]: 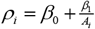 . The *β*_1_ parameter allows the risk attitude to vary by farm size.

Finally, we consider the following profit distributions for the two types of seeds

#### Conventional seed

Assume the profit distribution function of the conventional seed technology is known to farmers, and defined as 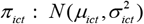, where *μ*_*ict*_ is the average profit perceived by farmer *i* at time *t* and 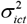 is its corresponding variance. Further, the variance may depend on a random state variable *z*_*t*_ which follows an AR(1) process. For example, random events such as pest infestation or weeds create profit uncertainty with potentially lasting effects. Then, we define the variance term as 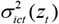 where *z*_*t*_ = λ *z*_*t*− 1_ + *v*_*t*_, and 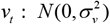 is the white noise added to the AR(1) process in each time period. We assume the random state variable brings additional uncertainty, thus 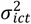 takes the form: 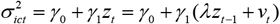, where *γ*_1_ > 0.

#### GM seed

The profit for the new technology is defined as *ρ* _*igt*_ = *μ*_*igt*_ + *ε* _*igt*_, where

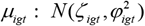 is the perceived average profit of GM seed for farmer *i* at time *t*, and 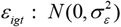is the error term following an i.i.d. normal distribution with zero mean and constant variance. It includes the impact of unpredictable weather, soil quality and unobserved farmer characteristics. Farmers form beliefs about the distribution of *μ*_*igt*_, which can be updated over time based on learning from own experience and interactions with neighbors.

### Farmer learning about GM seed profitability

We assume that farmers learn about the mean profitability of GM seed technology in a Bayesian fashion. Specifically, the information set of farmer *<u>i</u>* at time 0, *I*_*i*0_ consists of exogenous information (e.g., this information can come from agronomists or agricultural extension agents or from farmers’ own observations of pest and weed infestation in past years and possible effectiveness of GM seeds) on the GM average profit ζ _*ig*0_ and its accuracy 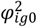. At time 1, farmer *i* may partially experiment with GM seeds and then, using information from his field experiments and/or from neighbors, updates the information set to *I*_*i*t_ with beliefs on both parameters ζ_*ig*1_ and 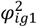. This process continues until learning is complete. As a note, our data do not include information on the actual profits, so we assume that farmers’ beliefs on GM mean profit ζ_*idt*_ is constant over time. Farmers receive an unbiased estimate of it initially, and then update their beliefs on its accuracy, 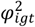 in later time periods according to the Bayesian rule as follows (see [22], Appendix A for detailed derivation):

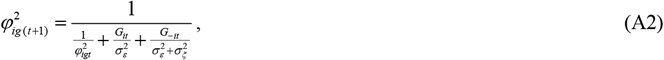

where *G*_*it*_ is the number of plots planted by farmer *i* with the GM seed at time t, *G*_−*it*_ is the average number of plots planted by his neighbors at time *t*, and 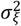 is the additional known variance or noise in learning from neighbors. The coefficients 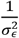 and 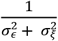 can be interpreted as the weights farmers attach to self and social information sources. In general information gained from own experience is more precise than that obtained from neighbors’ experience: 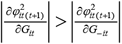

### Farmer adoption process

We assume heterogeneity among farmer *i*’s farmland due to differences in soil quality, infestation vulnerability, or other factors relating to agricultural production conditions. Farmer *i* can arrange his land in plots such that the suitability to plant GM seeds is decreasing in plot order. Let plots be ranked by *k*_*i*_ = 1,2, *A*_*i*_ from high to low suitability for GM seeds. Then, for each plot *k*_*i*_ the difference between the unbiased belief of GM seed mean profit 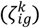 and the conventional seed mean profit 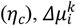, is

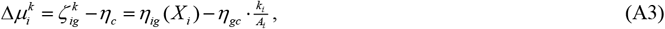

where *η*_*ig*_ is the upper bound of the profit difference, which is a function of farmer *i’s* characteristics *χ*_*i*_. Assume *η*_*gc*_ > 0 so that the mean profit difference between the two types of seeds is decreasing in *k*_*i*_.

Using equation (A3), if farmers’ adoption decisions are made based on comparing mean profits only (without forward-looking) and there is independence of profits from different land plots, the mean and variance of the total profit take the following form:

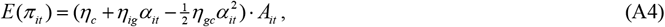

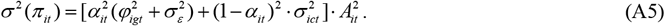

Then, the current payoff at time *t* for farmer *i* is:

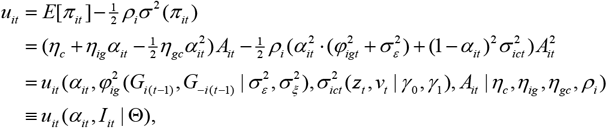

where the information set *I*_*it*_ consists of all factors that affect current expected utilities and/or probability distribution of future expected utilities. It includes current belief of GM profit variance 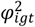, profit variance of conventional seeds 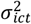, and total soybean acreage *A*_*it*_. The set Θ is the model’s parameter space, defined as

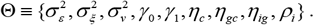

